# Expression of complexes I, II and IV in the SGNs of noise-stimulated rats was decreased, and mitochondrial energy metabolism was disturbed, mediating the damage and degeneration of SGNs

**DOI:** 10.1101/2022.08.19.504483

**Authors:** Zhong Jia Ding, Yin Wang, Ren Feng Wang, Wen Juan Mi, Jian Hua Qiu, Ding Jun Zha

## Abstract

Noise-induced hearing impairment can mediate delayed injury of spiral neurons (SGNs), resulting in degeneration of nerve fibers, synaptic degeneration and even death of SGNs. We believe that delayed injury is related to mitochondrial energy metabolism disorders. Therefore, we investigated ATP and the electron transport chain (ETC) in rat SGNs after noise injury and found that with prolonged injury time, ATP synthesis and the expression of complexes II and IV decreased, indicating the functional decline of the ETC. The maintenance of ETC function is related to subunit import and assembly of the complex. The disulfide relay mechanism controlled by the apoptosis inducing factor/coiled-coil-helix-coiled-coil-helix domain-containing protein 4(AIF/CHCHD4) pathway can regulate mitochondrial protein import. The results showed that AIF expression in SGNs decreased after noise exposure, indicating that noise damage to SGNs can restore intramitochondrial protein input by downregulating the AIF/CHCHD4 pathway, hindering ETC function; insufficient ETC function is a possible reason for the delayed injury of SGNs.

## Introduction

Mitochondria are the energy factories in cells, and the activities and degeneration of cells are closely related to the ATP produced by the mitochondria. Spiral neurons (SGNs) also degenerate during noise exposure-induced hearing loss. As early as 1988, a study found that SGN afferent fibers swelled under noise exposure(Puel and Ruel et al., 1998), and in 2018, in the adult mouse transient threshold shift model, SGNs exhibited acute loss and ribbon synaptic damage (Parthasarathy and Kujawa, 2018). Scholars have focused on the mitochondria-dominated apoptosis mechanism, and the relationship between these changes and energy metabolism disorders is not well understood.

ATP is the cellular energy currency, and the electron transport chain (ETC) is the main pathway by which mitochondria produce ATP. Changes in the complexes in the ETC will not only generate oxidative free radicals and other damaging factors that induce apoptosis but also reduce the efficiency of the redox system(Spinelli and Haigis, 2018). For example, complex I (ubiquinone oxidoreductase) in the ETC is not only the largest oxidizer in the ETC but also the rate-limiting enzyme of the entire respiratory movement (Sharma and Lu et al., 2009; Srinivas, 2017). Therefore, factors affecting the assembly and stability of complex I can lead to imbalances in energy metabolism, affect normal cell activities, and even induce cell degeneration. Likewise, changes in other complexes can affect the efficiency of ATP synthesis. However, the relationship between the structural and functional changes of SGNs and mitochondrial energy metabolism during noise exposure is less clear, and the structure of SGNs will continue to change for a period of time after the noise is stopped. Without considering the stimulation of damage factors, we suspect that energy metabolism disorders may be related to the structure of SGNs. Therefore, we aimed to disable SGNs through noise, observe the structural changes of SGNs and the changes in the ETC complex and ATP levels, and explore the possible related mechanism of noise damage to the ETC to provide a new research direction for the protection of SGNs.

## Result

### 1. High-intensity noise leads to an increased shift in hearing threshold and a decreased ABR I wave amplitude

We exposed adult SD rats (180 g) to noise stimulation (stimulation conditions: 110 dB SPL, 2 hours a day, costimulation for 7 days) and observed the ABR threshold on the 1st, 3rd, 5th, and 7th days. We found that the ABR threshold significantly increased and reached a stable level on Day 7: click, 50.00±10.48 dB; 4 kHz, 49.16±9.17 dB; 8 kHz, 49.16±9.17 dB; 16 kHz, 50.00±8.36 dB; 32 kHz, 60.00±6.32 dB. The new threshold was 30.16±8.15 dB higher than the normal ABR threshold (P<0.05), which indicated permanent hearing damage in the rats. The damaging effect on auditory afferent primary neurons (SGNs) was verified by detecting the amplitude of the ABR I wave. It was found that the amplitude of the ABR I wave decreased under noise stimulation at various decibel levels. After the stimulation ended, the amplitude of the ABR I wave rebounded, and there was no significant difference compared with the amplitude before the noise stimulation. As shown in Figure 1, the existence of residual hearing was verified. However, the amplitude of the ABR I wave decreased significantly during 40 dB and 50 dB noise stimulation, suggesting that the spiral neurons in the auditory conduction pathway were damaged.

**Figure 1.**
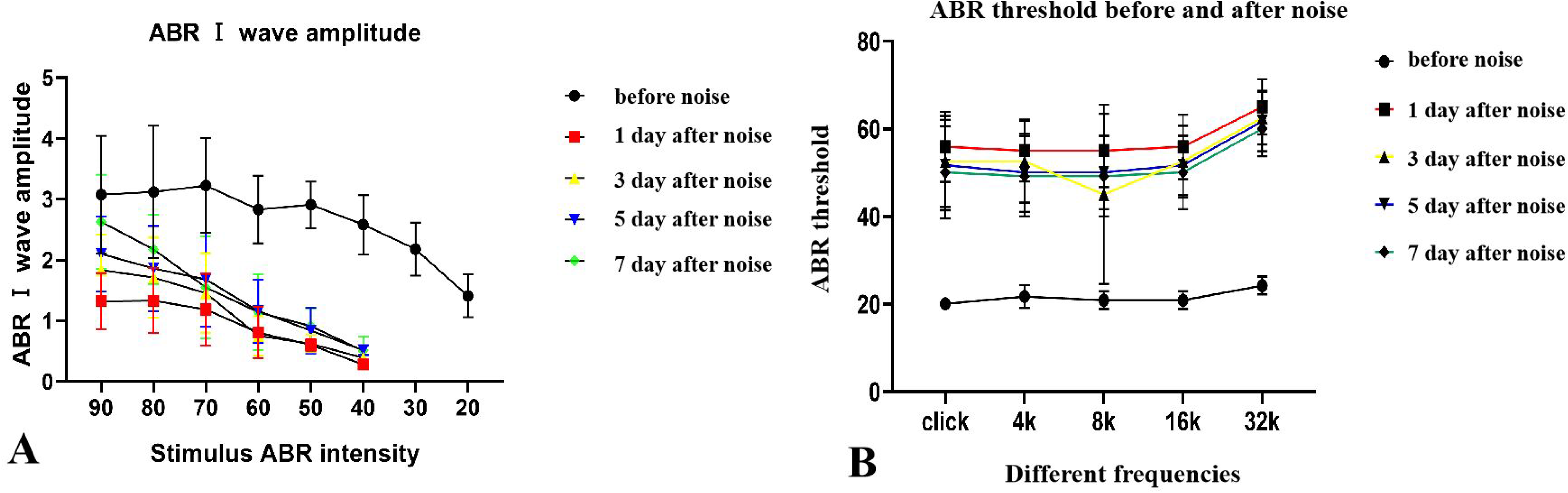
Audiogram of rat after noise exposure. A represents the waveform of the amplitude of the ABR I wave before and 7 days after noise exposure, which can reflect the health status of SGNs in the auditory pathway. The amplitude of the ABR I wave is essentially stable 7 days after noise exposure. Acoustic stimulation of 90 dB caused a higher amplitude, while the amplitude significantly decreased after 40 dB acoustic stimulation. With an increase in the sound stimulus dB level, SGNs become more responsive to sound. B represents the waveform of the ABR threshold before and 7 days after noise exposure. The ABR threshold is essentially stable 7 days after noise exposure, and the threshold is 30.16±8.15 dB higher than the normal threshold in each frequency band.

### 2. Noise causes helical neuron loss and postsynaptic AMPAR decline

To verify the effect of noise on rat helical neurons, we divided the rat cochlea into axial serial frozen sections and used β-tubulin (green) to identify type I spiral ganglion neuron cell bodies, GluR2A (red) to identify AMPARs, and DAPI (blue) to identify the nuclei. All the spiral neuron cell bodies in the serial frozen sections were manually counted, and the differences were compared. It was found that the number of spiral neurons was indeed significantly reduced after noise exposure, as shown in Figure 2. The red GluR2A immunofluorescence was semiquantitative and showed that the expression of AMPARs decreased, as shown in Figure 3. Similar conclusions were obtained by using western blotting to verify the decreased expression of GluR2A, which was positively correlated with the reduction in the number of spiral neurons, as shown in Figure 4, indicating that noise stimulation can effectively reduce synapses. The expression of back-end AMPARs mediates the loss of spiral neurons, and we consider that this mechanism may be related to the imbalance in mitochondrial energy metabolism.

**Figure 2.**
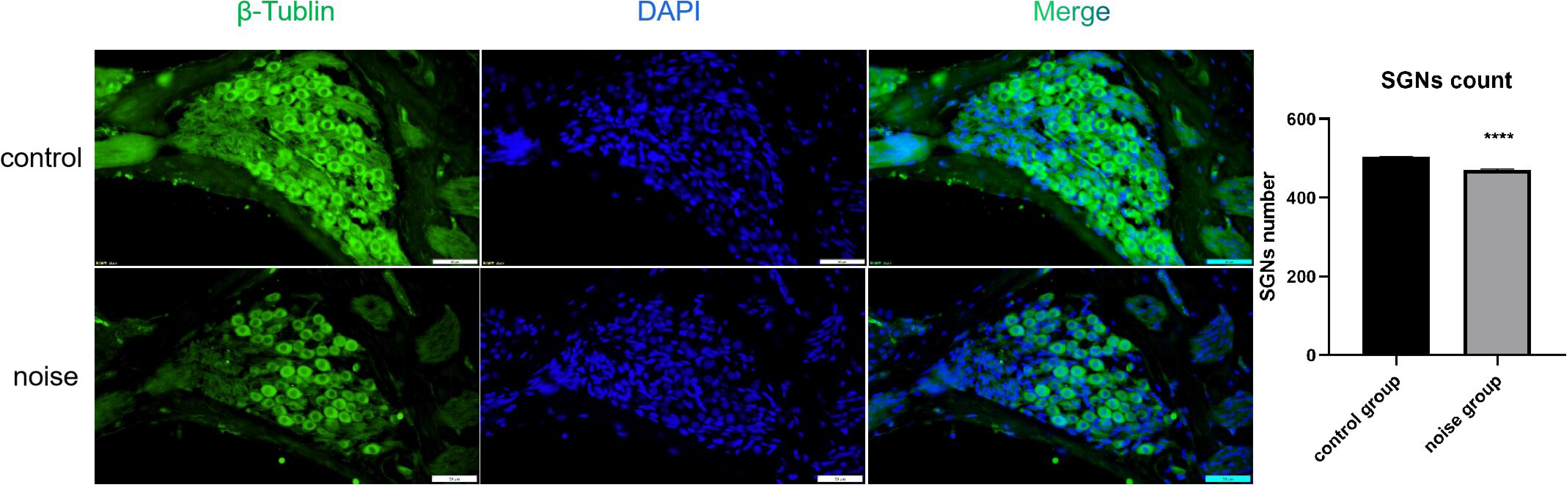
Immunofluorescence and count of SGNs in the spiral ganglia before and after noise exposure. β-Tubulin (green) stains intracellular nerve fibers in SGNs, and DAPI (blue) stains the nuclei; the merged image is a combination of both stains. We manually counted the number of cell bodies in frozen sections of all SGNs, and the histogram indicated that the number of SGNs decreased after noise exposure compared with that of the normal group (P<0.05).

**Figure 3.**
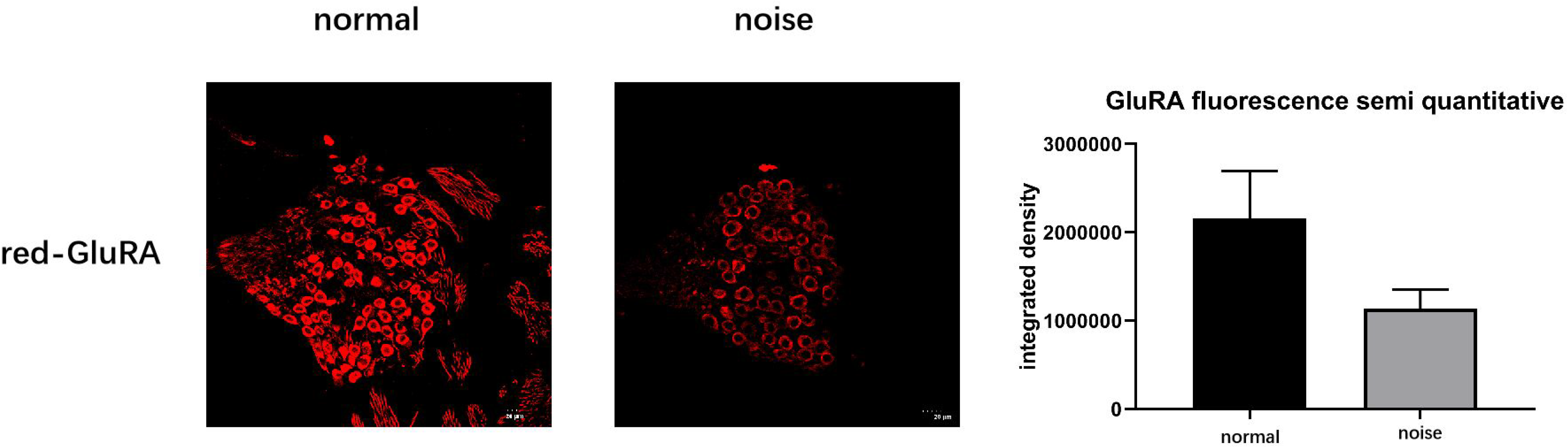
Fluorescence semiquantitative plots of GluRA in SGNs before and after noise exposure. Red color indicates the presence of GluR2A antibody at the spiral ganglion, which is a part of AMPARs in SGNs. Under the same fluorescence intensity exposure, the expression of AMPARs before and after semiquantitative noise exposure was found to decrease, as shown in the histogram, indicating that noise affects postsynaptic receptors in SGNs in a destructive manner.

**Figure 4.**
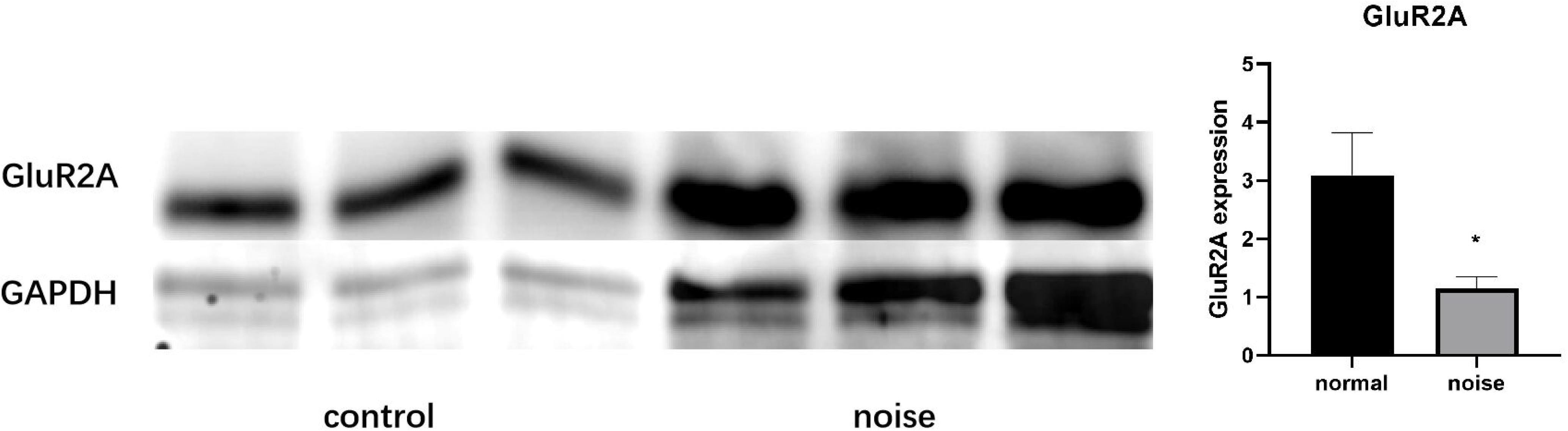
WB expression of GluR2A before and after noise exposure. By grayscale map and grayscale analysis, it was found that noise exposure reduces the expression of GluR2A and has a detrimental effect on postsynaptic AMPARs in SGNs.

### 3. Noise can lead to an imbalance in electron transport chain complexes and a decrease in mitochondrial ATP synthesis

Considering that most cellular energy is generated by the mitochondrial electron transport chain, we used an antibody against the mitochondrial complex to detect the mitochondrial ETC complex in SGNs before and after noise exposure by WB. As shown in Figure 5, the decrease in the expression of complexes II and IV was statistically significant, and the decrease in complex I expression was a trend only. The expression changes in ND3 and ND6 were detected by WB, and it was confirmed that the expression of ND subunits decreased. This decrease indicated that the function of complex I was affected. To directly verify the energy disorder of spiral neurons after noise exposure, we examined the ATP synthesis in the neurons and found reduction compared to the normal configuration. The resulting decrease in energy consumption, as shown in Figure 6, indicates that the damaging effect of noise on helical neurons may occur due to mitochondrial energy metabolism, and the imbalance of energy metabolism characterized by damage to electron transport chain complexes restricts the transmission of postsynaptic signals and mediates spiral neuron damage.

**Figure 5.**
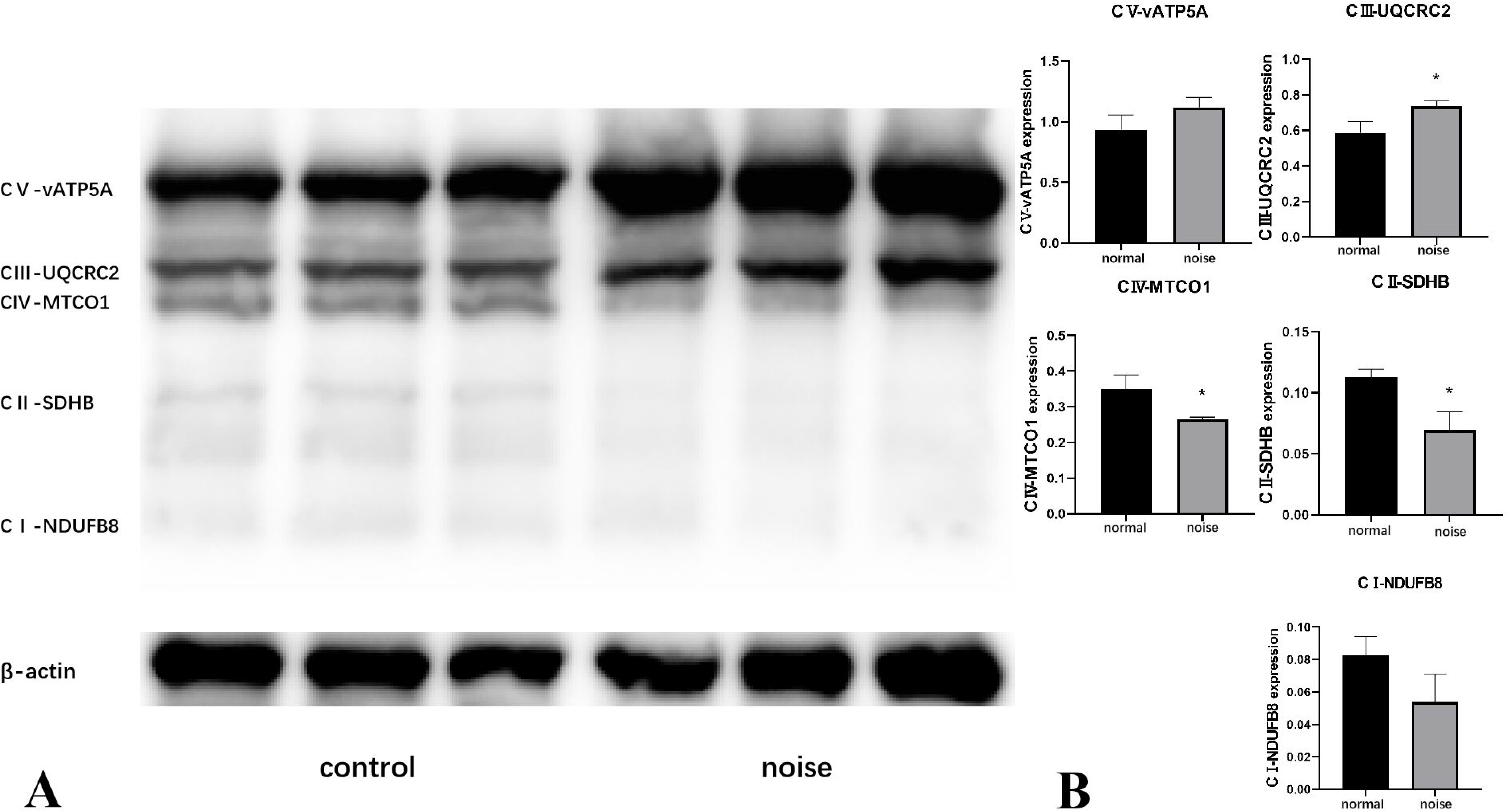
Expression of mitochondrial complexes before and after noise exposure. A is the grayscale image of the mitochondrial complex before and after noise exposure. From top to bottom, the bands show the subunit vATP5A of complex V, the UQCRC2 subunit of complex III, the MTCO1 subunit of complex IV, the SDHB subunit of complex II, and the NDUFB8 subunit of complex I. β-actin was used as the positive control. B is the quantitative histogram of the grayscale analysis. The expression of complex V and complex III increases after noise exposure, while the expression of complex I, II, and IV decreases. Only the decrease in the expression of complex II and complex IV was significant (P<0.05).

**Figure 6.**
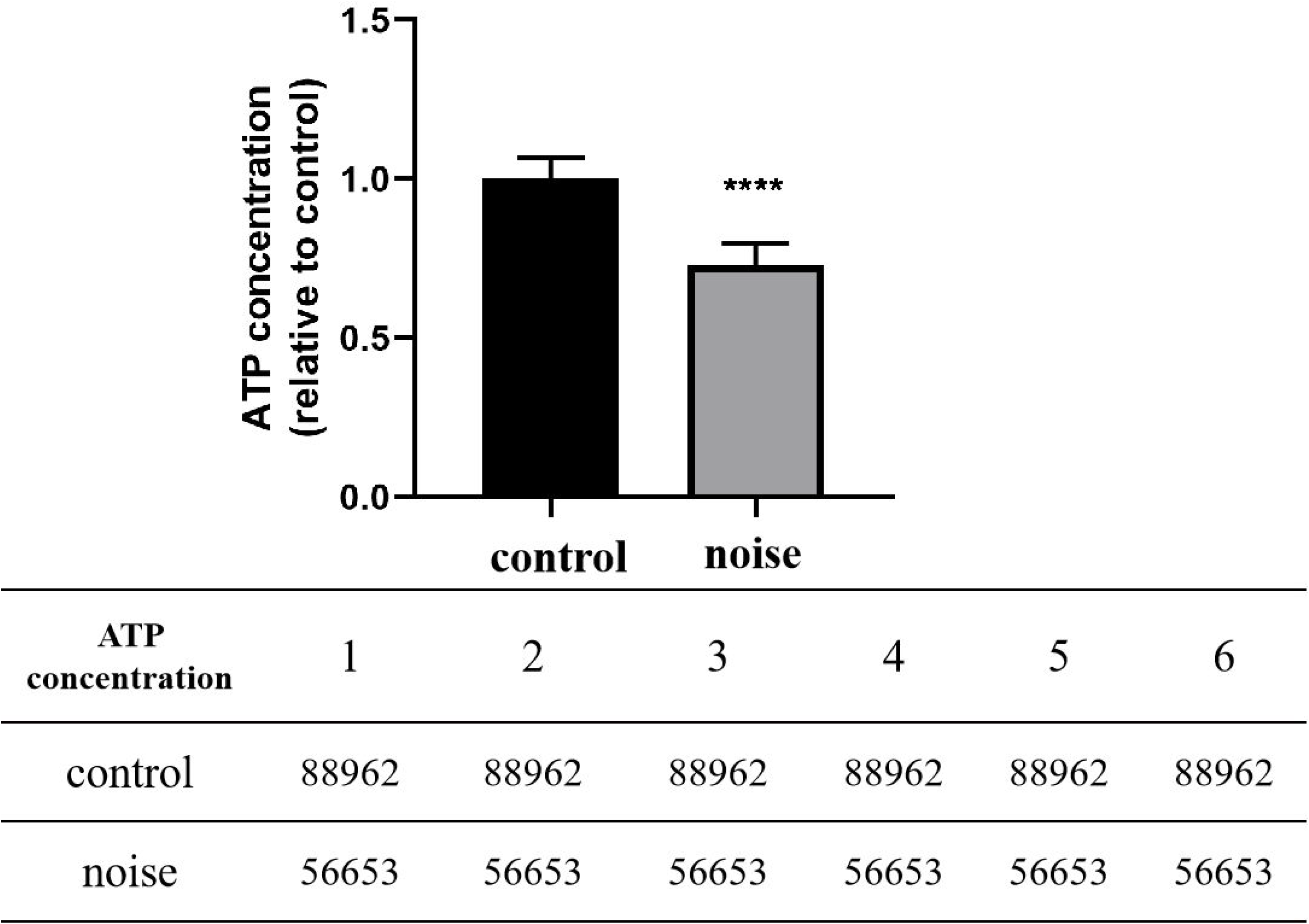
Determination of relative amounts of ATP synthesis before and after noise exposure. The control group was the ATP content in normal SGNs, and the intervention group was the ATP content in SGNs after noise exposure. As shown in the figure, the synthesis of ATP after noise intervention was significantly lower than that in the control group (P<0.05).

### 4. Decreased expression of AIF (AIF) may be related to complex dysfunction

Mitochondrial complexes are located in the inner mitochondrial membrane and matrix, but the formation of complex subunits is mostly performed by nuclear DNA. Therefore, the passage of subunits into the mitochondrial matrix is a key pathway that restricts the synthesis, assembly and functional repair of complexes. It is currently known that the disulfide relay mechanism controlled by the AIF/CHCHD4 pathway can regulate mitochondrial protein import, including complex I, and maintain stable mitochondrial functions. After noise exposure, we found that the expression of AIF in helical neurons decreased, as shown in Figure 7, which may impair the assembly and function of mitochondrial complexes.

**Figure 7.**
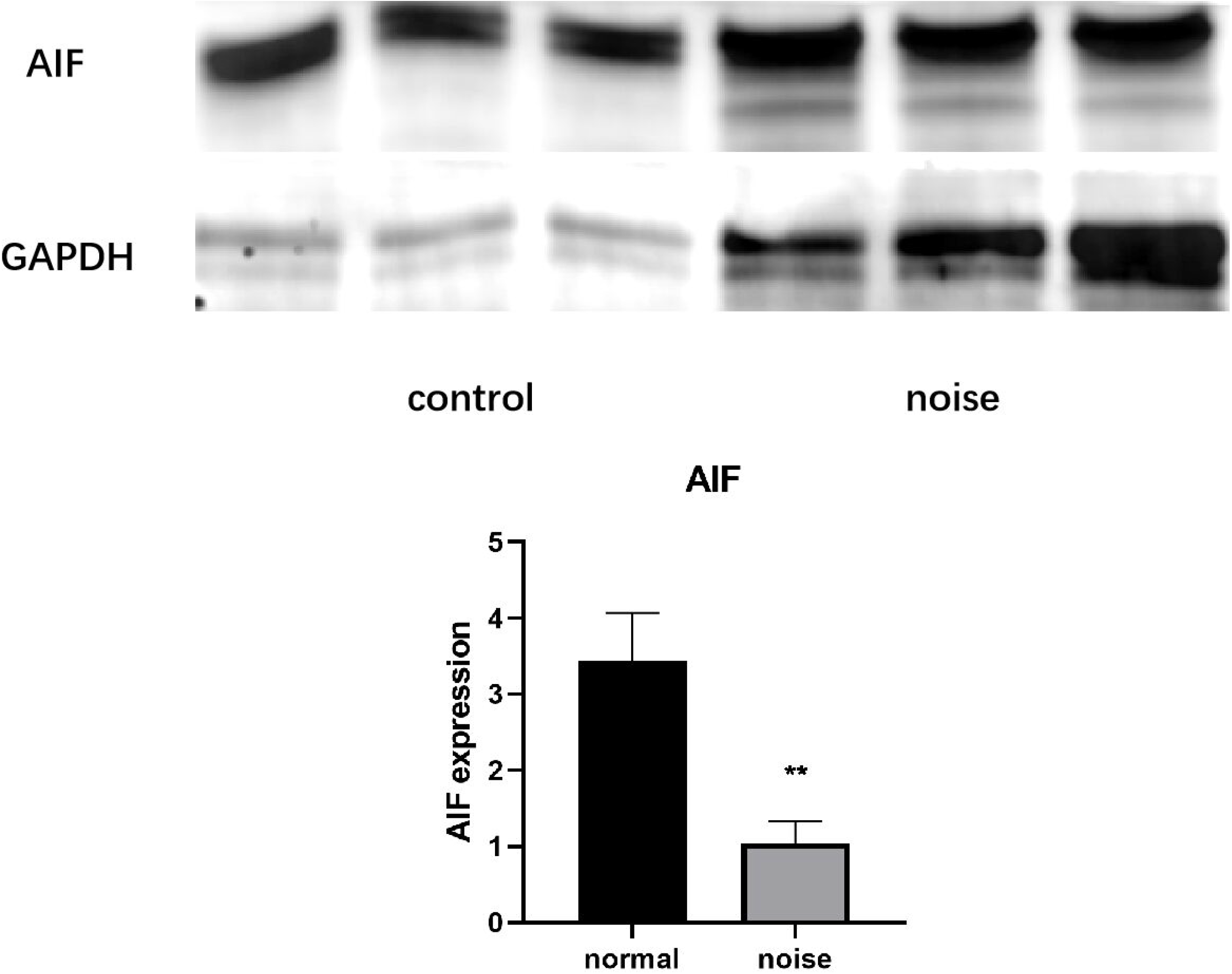
Grayscale images of AIF expression before and after noise exposure. The upper picture is the grayscale image of AIF and housekeeping gene GAPDH expression in western blot. The three bands on the left are from the control group, and the three bands on the right are from the noise group. The lower picture is the grayscale analysis histogram, which shows that AIF expression after noise exposure was significantly lower than normal (P<0.05).

## Discussion

Noise can cause metabolic disorders in the inner ear cells and affect hearing. In addition to hair cells, SGNs are also affected (Liberman and Kujawa, 2017). However, after the noise stops, the structure of SGNs will continue to change, including afferent fiber degeneration and reduction of ribbon synapses, and the number of SGN cells will decrease(Wang and Ren, 2012). We suggest that these changes are closely related to an energy metabolism disorder in SGNs. The energy metabolism of SGNs is inseparable from the oxidative phosphorylation process of mitochondria. Mitochondria are the core of cellular energy metabolism and the key to energy production in eukaryotic cells. The Krebs cycle transfers energy to the carriers NADH and FADH, while oxidative phosphorylation (OXPHOS) can use the proton gradient to redox NADH and FADH and finally synthesize the cellular currency ATP to supply the functional needs of SGNs(Xu and Xue et al., 2020). We detected a significant decrease in ATP synthesis in SGNs 1 week after the noise stimulation stopped, indicating that although the noise stimulation was removed, the damaged SGNs continued to degenerate, which may be related to a lack of energy required to maintain a normal state, thus showing continued degradation from synapse to soma. Oxidative phosphorylation is the main process of mitochondrial ATP synthesis, carried out by the electron transport chain (ETC), which consists of four polymerases, two mobile electron carriers, and one ATP synthase (complex V). The four polymerases include complex I (NADH: CoQ oxidoreductase), complex II (succinate: ubiquinone oxidoreductase), complex III (coenzyme Q: cytochrome c oxidoreductase), and complex IV (cytochrome c reductase), and the two electron carriers are NADH and FADH (Signes and Fernandez-Vizarra, 2018). Respiration transfers H+ to NADH and FADH through the action of the complex, forms a proton gradient to condense ADP and inorganic phosphate, combines with oxygen to form water, transfers electrons from the carrier and releases energy to supply ATP synthesis. Among the complexes, complex I is the key to the formation of a proton gradient and the rate-limiting step for the carrier to enter the respiratory chain in the ETC(Fiedorczuk and Sazanov, 2018), while complex V synthesizes ATP from ADP and inorganic phosphate as dictated by the energy supply. ATP deficiency is closely associated with aging, genomic instability, apoptosis, inflammation, and neurodegenerative diseases (Picca and Calvani et al., 2021). We performed ETC complex detection on SGNs after noise exposure and found that the expression of complex I, complex II, and complex IV and the synthesis of ATP decreased, indicating that mitochondrial energy metabolism was damaged by noise exposure. In addition, the number of AMPARs declined in SGNs. The decrease in receptors and the number of cells may also be caused by the degeneration of SGN cells due to insufficient energy. Therefore, noise damage affects the energy metabolism of SGNs and leads to their degeneration and apoptosis, which continue after noise stops due to the continuous decline in energy supply. However, how noise regulates the changes in mitochondrial complexes that cause energy metabolism disorders remains to be further studied.

Scholars believe that mitochondria are the main organelles for the production of oxidative free radicals in the body, which are the main cause of cell damage and apoptosis(Johnson and Mercado-Ayon et al., 2021). However, we found that for a period of time after the noise stopped, the expression of the mitochondrial complex decreased, the working efficiency of the ETC was reduced, and there was no continuous harmful stimulus to mediate the generation of free radicals. Therefore, determining the reason for the degeneration of SGNs due to the energy metabolism disorder is beneficial. The complication is mainly caused by the decline in the expression of the important components of the ETC, complex I, complex II, and complex IV. Recent studies have found that the mitochondrial complex is in a state of dynamic equilibrium and needs continuous input from the nucleus for gene translation and expression through the AIF/CHCHD4 mitochondrial disulfide relay system to mitigate the high loss of the subunit in the proton gradient(Reinhardt and Arena et al., 2020). AIF (apoptosis-inducing factor) can mediate apoptosis (Herrmann and Riemer, 2021), but it normally acts as a flavoprotein in the mitochondrial matrix to complete the relay system input. AIF deletion directly mediates the decline in complex I function. In AIF conditional knockout mice, AIF deficiency leads to the suspension of the P module in complex I and the accumulation of the Q module, which affects the function of complex I(Troulinaki and Buttner et al., 2018; Zong and Zhao et al., 2020). There are also studies confirming that the expression of complex III and complex IV is affected(Cabon and Bertaux et al., 2018). We found that the expression of AIF decreased significantly in SGNs after noise injury, indicating that the AIF/CHCHD4 input system was affected, which may be related to the decreased expression of complex I, complex II, and complex IV. Therefore, timely regulation of ATP synthesis can reverse mitochondrial energy metabolic disorders and protect auditory cell SGNs, which provides a new direction for research into the continuous degeneration of noise-induced deafness.

## Methods

### 1. Animals

All experiments were performed on Sprague Dawley rats (200 g, 8-9 weeks) obtained from the Laboratory Animal Center of Fourth Military Medical University (FMMU), Xi’an Shaanxi Province, China. The animals were maintained under standard laboratory conditions (12 h dark/light cycle, temperature 22–26 °C, air humidity 40– 60%) with food and water available ad libitum. The experimental protocols were approved by the Institutional Animal Care and Use Committee of FMMU. The rats were randomly divided into two groups (n = 10 per group): ① Control group, experimentally naive rats that were exposed to ambient sound levels (50-60 dB SPL, measured with a sound level meter) without noise exposure; ② Noise group, rats that were exposed to intense noise. Hearing tests were performed on Days 1, 3, 5 and 7, and the rats were anesthetized and killed on day

### 2. Noise exposure

The noise exposure was conducted in a noise sound box. The animals were placed in metal wire cages with free access to food and water for one week to acclimate to the noise sound box. During noise exposure, water bottles were located on the side of the cages, and there were no objects between the speakers and the rats’ ears. The noise (white noise, 110 dB SPL) was generated by a RadioShack Supertweeter located above the cages and was amplified with a power amplifier (Yamaha AX-500U, Japan) and a loudspeaker. The noise level was monitored with a sound level meter (Bruel and Kjaer, Typer 2606), with a variation less than 5 dB across the space available to animals. The animals were exposed to noise 2 h per day for 7 days.

### 3. Auditory brainstem response (ABR)

The ABR thresholds in rats were measured at 1 day, 3 days, 5 days and 7 days after noise exposure. Under light anesthesia with 10% chloral hydrate (400 mg/kg), the active, reference and ground needle electrodes were inserted beneath the skin at the vertex, the mastoid area of the test ear and the contralateral mastoid, respectively. The TDT III auditory-evoked potential workstation was used for sound generation and presentation, and data acquisition was performed with SigGenRZ and BioSigRZ software (Tucker Davis Technologies, Fort Lauderdale, FL, USA). The ABR was elicited by tone bursts (8, 16, 32 kHz; 0.5-millisecond rise/fall time, no plateau, alternating phase) or broadband clicks (10 milliseconds) presented at 21.97 s−1. The stimulus was played through a high-frequency speaker (model: MF1 Multi-Field Magnetic Speakers) located approximately 2 cm in front of the test ear. The intensity of the stimulus was decreased in 5-dB steps until the evoked responses disappeared. The differential potential was sampled over 10 milliseconds, filtered (low pass, 4 kHz; high pass, 100 Hz) and averaged (512 sweeps of alternated stimulus polarity) to obtain mean traces at each intensity(Liu and Lu et al., 2019). The lowest intensity that was able to elicit a two-phase waveform from 5 to 15 ms after signal onset was considered the ABR threshold.

### 4. Immunofluorescence staining

Slips of SGNs were fixed in 4% paraformaldehyde for 30 min and permeabilized in 0.1% Triton X-100 for 15 min. After washing with PBS, the samples were incubated in a blocking solution of bovine serum albumin (BSA, 5%, Sigma, USA) for 20 min, followed by incubation with antibodies against AIF (1:200, rabbit, Abcam, USA) and β-tubulin (1:200, mouse, Abcam, USA) for 24 h overnight at 4 °C. Alexa 488-conjugated goat anti-rabbit (1:200, Invitrogen, USA) and Alexa 594-conjugated donkey anti-mouse (1:200, Invitrogen, USA) were used to label the primary antibodies by incubation for 40 min at 37 °C. Stained SGNs were observed under a fluorescence microscope (Olympus, Japan), and three photos of different groups were taken by the microscope.

### 5. Mitochondrial isolation and purification

Neurons were lysed with a lysis buffer containing protease inhibitors. The cell lysate was centrifuged for 10 min at 750 g at 4 ℃, and the pellets containing the nuclei and unbroken cells were discarded. The supernatant was then centrifuged at 15 000 × g for 15 min. The resulting supernatant was removed and used as the cytosolic fraction. The pellet fraction containing the mitochondria was further incubated with PBS containing 0.5% Triton X-100 for 10 min at 4 °C. After centrifugation at 16 000 × g for 10 min, the supernatant was collected as the mitochondrial fraction.

### 6. Measurement of ATP synthesis

A luciferase/luciferin-based system was used to measure ATP synthesis in isolated mitochondria as described elsewhere (Parone et al., 2013). Thirty milligrams of mitochondria-enriched pellets was resuspended in 100 ml of buffer A (150 mM KCl, 25 mM Tris HCl, 2 mM potassium phosphate, 0.1 mM MgCl2, pH 7.4) with 0.1% BSA, 1 mM malate, and 1 mM glutamate and buffer B (containing 0.8 mM luciferin and 20 mg/ml luciferase in 0.5 M Tris-acetate pH 7.75). The reaction was initiated by the addition of 0.1 mM ADP and monitored for 120 min using a microplate reader at 530 nm. There were six samples in each group, and the experiment was repeated a minimum of 3 times.

### 7. Western blot analysis

Total protein concentrations were measured using the Pierce BCA method (Sigma, USA). Equivalent amounts of protein (40 mg per lane) were loaded and separated on 10% SDS-PAGE gels and transferred to polyvinylidene difluoride (PVDF) membranes. Membranes were blocked with 5% skim milk solution in Tris-buffered saline with 0.1% Triton X-100 (TBST) for 1 h and then incubated overnight at 4 ℃ with primary AIF OXPH, complex I ND1, ND3, ND6, GluRA, or b-actin (1:500, #4970, Cell Signaling, USA) antibody dilutions in TBST. Then, the membranes were washed and incubated with secondary antibodies (Santa Cruz, USA) for 1 h at room temperature. Immunoreactivity was detected with Super Signal West Pico Chemiluminescent Substrate (Thermo Scientific, Rockford, IL, USA). ImageJ (Scion Corporation) was used to quantify the optical density of each band. The expression of each protein was calculated from the optical density of each band normalized against the optical density of b-actin and expressed as the fold change compared to the control levels.

